# SMARCA4 is essential for early-stage tumor development but its loss promotes late-stage cancer progression in small-cell lung cancer

**DOI:** 10.1101/2025.05.21.655382

**Authors:** Nicole A. Kirk, Jin Ng, Kate-Lin Ly, Young Ho Ban, Godfrey Dzhivhuho, Jinho Jang, Kyung-Pil Ko, Michael S. Kareta, Jae-Il Park, Anthony N. Karnezis, Anish Thomas, Kate D. Sutherland, Kwon-Sik Park

**Affiliations:** Department of Microbiology, Immunology, and Cancer Biology, University of Virginia, Charlottesville, Virginia, USA; ACRF Cancer Biology and Stem Cells Division, The Walter and Eliza Hall Institute of Medical Research, 1G Royal Parade, Parkville, Victoria, Australia; Department of Medical Biology, The University of Melbourne, Parkville, Victoria, Australia; Cancer Biology and Immunotherapies Group, Sanford Research, Sioux Falls, South Dakota, USA; Department of Experimental Radiation Oncology, MD Anderson Cancer Center, Houston, Texas, USA; Department of Pathology and Laboratory Medicine, University of California Davis, Sacramento, California, USA; Developmental Therapeutics Branch, Center for Cancer Research, National Cancer Institute, Bethesda, Maryland, USA

**Keywords:** SMARCA4, SCLC, NE differentiation

## Abstract

SMARCA4 and other components of the SWI/SNF chromatin remodeling complex have been implicated in various cancers. Yet, its role in small cell lung cancer (SCLC) tumorigenesis remains poorly understood. Genetically engineered mouse models (GEMMs) of SCLC revealed that deletion of *Smarca4* significantly decreased tumor development in this model. Pharmacological inhibition of SMARCA4 decreased the proliferation of preneoplastic neuroendocrine (NE) cells. These effects coincided with reduced expression of the lineage-specific transcription factor, ASCL1, suggesting that disruption of the SMARCA4-ASCL1 axis impairs tumor development. However, *Smarca4*-deficient tumors, albeit smaller than controls, displayed features associated with malignant progression, including variant histology and the loss of NE differentiation. This prompted us to test the functional role of SMARCA4 in established tumor cells that recapitulate late-stage disease. Intriguingly, whilst *Smarca4* knockdown in tumor cells failed to affect their proliferative capacity *in vitro, Smarca4* knockdown tumors exhibited enhanced growth following subcutaneous transplantation in athymic nude mice. Interestingly, SMARCA4 knockdown significantly reduced expression and cell-surface display of PVR, a ligand for activating natural killer (NK) cells. These results led to an idea that the enhanced tumor formation was partly owing to altered tumor-NK cell interactions mediated by the SMARCA4-PVR axis in tumor cells. These findings suggest that SMARCA4 plays a temporally distinct role in SCLC, supporting early tumorigenesis but potentially functioning as a tumor suppressor in the later stages. The dramatic differences observed when targeting SMARCA4 in distinct disease states emphasize a need to acknowledge how differences in the timing of alterations can drastically alter tumor evolution.

## Introduction

SCLC accounts for 13% of all lung cancers and is an aggressive disease with a 5-year survival rate of 7% in patients(1). SCLC generally lacks actionable driver mutations such as mutant KRAS and EGFR found in NSCLC(2). Instead, it harbors intractable loss-of-function mutations in *RB1* and *TP53* and other tumor suppressor genes(2), presenting significant challenges in developing targeted therapies. Functional understanding of these alterations is necessary for gaining important insights into the biology of SCLC and to uncover novel preventive and therapeutic strategies.

Emerging evidence points to the SWI/SNF chromatin remodeling complex as a central player in cancer biology(3). The SWI/SNF complex controls gene accessibility to transcriptional machinery and its subunits, including the catalytic component and ATP-dependent helicase (SMARCA4), are mutated in around 20% of all cancers(4). SMARCA4 has been shown to act as a tumor suppressor in mouse models of lung adenocarcinoma(5,6), while its role in SCLC development remains largely unknown. Recent studies showed that pharmacological inhibition of SMARCA4 reduces the proliferation of SCLC cell lines and patient-derived cells(7–9) and knockdown of SMARCA4 causes vulnerability within cell lines deficient for MAX, a member of MYC family proteins(10). While indicating SMARCA4 dependency in SCLC cells, these studies relied on established tumor cell lines and have not directly addressed its role in SCLC development. It also remains unclear whether the inhibitory effect on some SCLC cells results from the dual-inhibition of SMARCA4 and SMARCA2, which may cause a synthetic lethality among the components of the SWI/SNF complex(11). While another study suggested that *Smarca4* knockout decreases tumor incidence in *Rb1*/*Trp53-*mutant GEMM of SCLC(12), the exact role of *Smarca4* in SCLC development remains unclear due to the scarce description of phenotypes in the GEMM and absence of mechanistic insight underlying the decreased tumor number. Whilst SMARCA4 mutations have been observed in a “real world” SCLC patient cohort(13), caution should be exercised as inactivating mutations in *SMARCA4* are associated with SMARCA4-deficient undifferentiated tumors (SMARCA4-UT), a newly described tumor that is a mimic of SCLC(14). These observations together with the recent reclassification of YAP1-positive SCLC human cell lines as SMARCA4-UT(15) contribute to the confusion around a role of SMARCA4 in SCLC. Despite the growing interest in exploring the SWI/SNF complex components as prognostic biomarkers and therapeutic targets, critical gaps exist in understanding the roles they play in SCLC.

To address these unresolved questions, we used GEMMs and complementary *in vitro* models to dissect the functional role of SMARCA4 across stages of SCLC development. Our findings reveal that SMARCA4 supports early tumor formation but may restrain tumor progression at later stages, highlighting a temporally distinct and dual role in SCLC biology. More broadly, this study underscores the complex interplay between chromatin remodeling, lineage plasticity, and immune evasion in the evolution of aggressive cancers.

## Results and Discussion

To determine the impact of losing SMARCA4 on SCLC development, we utilized a GEMM known as RPR2 mice. In this model, tumors are initiated by adenovirus Cre-mediated deletion of *Rb1, Trp53*, and *Rbl2* floxed alleles in the airway epithelium and recapitulate ASCL1-dominant human SCLC(16). Following genetic crosses, we generated *RPR2* mice and their littermates carrying additional homozygous floxed alleles of *Smarca4* (hereafter *RPR2S*) and infected them with adenovirus Cre via intratracheal instillation (**Figure 1A**). Six months after infection, lung tumor burden (tumor area/total lung area) was significantly lower in *RPR2S* mice compared to *RPR2* littermates (**Figure 1B**). *RPR2S* tumors had a significant decrease in the levels of proliferation marker phosphorylated histone H3 (pHH3) and concomitant increase in the levels of apoptotic marker cleaved Caspase 3 (CASP3) compared to controls (**Figure 1C, Supplementary Figure 1A**). Soft agar assays showed primary cells isolated from *RPR2S* tumors formed significantly fewer colonies than *RPR2* cells (**Figure 1D**), indicating the inhibitory effect of losing SMARCA4 on cell expansion *in vitro*. To determine the role of SMARCA4 in the initial stages of tumor development, we inhibited the protein in precancerous cells (preSCs), immortalized *Rb1*/*Trp53*-mutant NE cells derived from early lesions developed in GEMMs(17). Treatment of a dual SMARCA4/2 inhibitor, FHD-286, resulted in significantly fewer colonies than the vehicle treated cells in soft agar assays (**Figure 1E**). Together, these results suggest that SMARCA4 is necessary for SCLC development and its functional loss hinders the proliferative capacity of *RPR2* tumor cells and preneoplasia.

**Figure 1.**
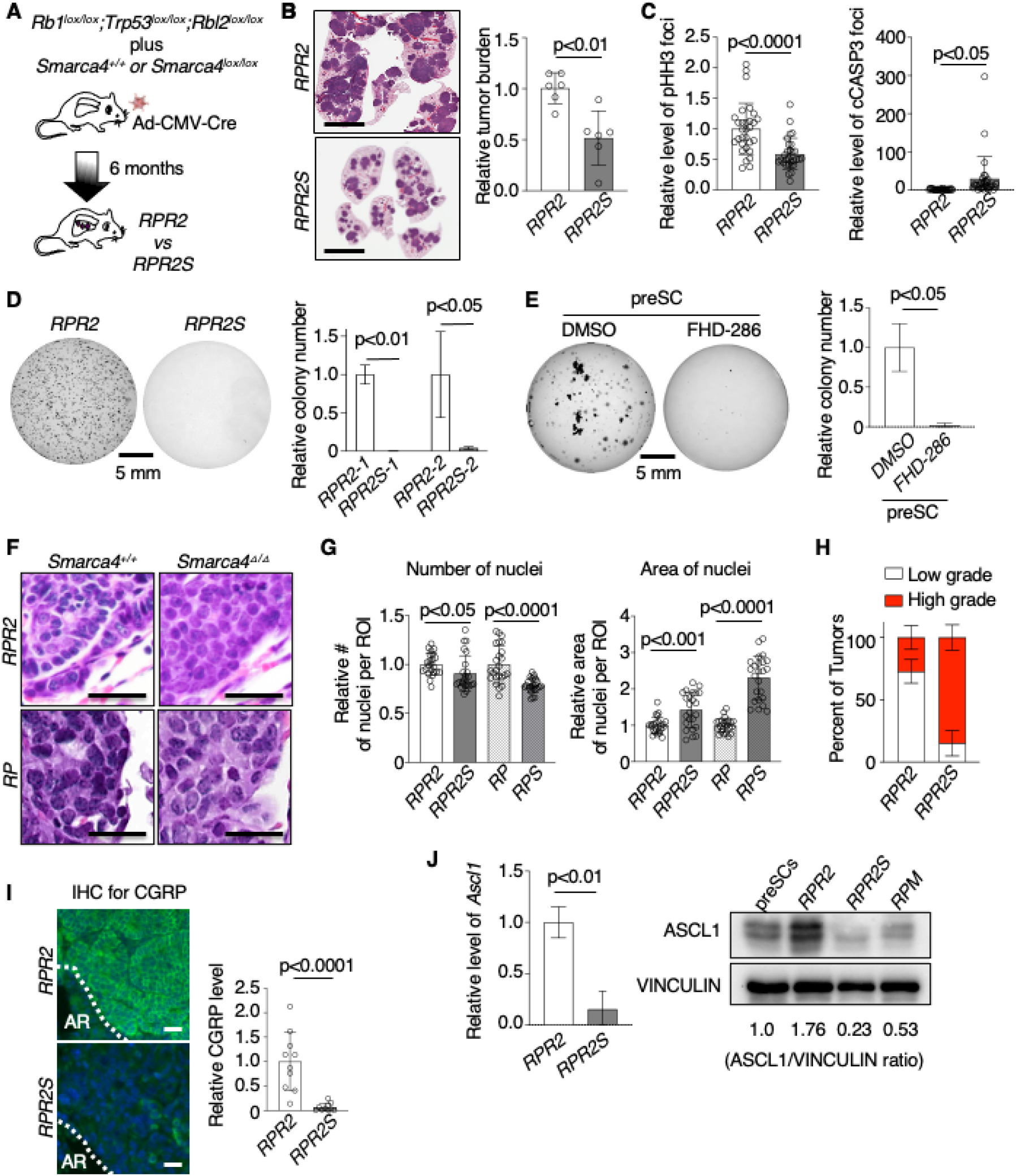
*Smarca4* loss inhibits tumor development in *RPR2* mice. **A**. Schematic of tumor induction in *RPR2* GEMM. **B**. Representative images of H&E-stained lung sections (scale bars: 5mm) and quantification of tumor burden in the GEMMs normalized to the control *RPR2* lungs (n=6 per group). **C**. Quantification of pHH3 foci and cleaved CASP3 foci within region of interest (ROI) in the lungs of *RPR2* and *RPR2S* (n=30 ROIs). **D**. Representative images of soft agar containing cells isolated from *RPR2* or *RPR2S* tumors and quantification of colony number. **E**. Representative images of soft agar containing preSCs treated with either DMSO or FHD-286 and quantification of colony number (n=3). **F**. Representative H&E images of the sections of lungs in GEMMs as indicated. Scale bars: 50µm. **G**. CellProfiler quantification of average number of nuclei and average size of nuclei per ROI from H&E-stained slides (**F**) (n=24 ROIs). **H**. Plots of percentage of low-grade and high-grade tumors present in the lungs of *RPR2* (n=7) and *RPR2S* (n=6) mice. **I**. Representative images of CGRP-stained lung sections. Scale bars: 50µm, and quantification of CGRP stain (n=10 per group). AR: alveolar region. **J**. RT-qPCR data showing *Ascl1* transcript level, normalized to levels of *Gapdh*, in *RPR2* and *RPR2S* cells (**D**) (n=3 per group) and immunoblot of ASCL1 and VINCULIN proteins isolated from various cells as indicated. Ratio of ASCL1 levels relative to VINCULIN levels, measured by densitometry, shown on the bottom.

To elucidate the mechanism by which *Smarca4* loss inhibits tumor development, we examined the histological features of *Smarca4*-deficient tumors. Unlike the classical NE morphology observed in RPR2 tumors and *Rb1/Trp53*-deficient tumors (henceforth *RP*)(18) (**Figure 1F**) – characterized by scant cytoplasm and rosette-like structures – RPR2S and *Smarca4*-deficient RP (RPS) tumors displayed variant histology, including increased cytoplasmic volume and distinct cell boundaries (**Figure 1F, Supplementary Figure 1B**). These differences were measured using CellProfiler to confirm an increase in nuclei size and decrease in nuclei number per ROI in the *Smarca4*-deficient tumors (**Figure 1G**). Classification by an independent pathologist revealed that there was an increase in the proportion of tumors classified as high-grade NE carcinoma in the *RPR2S* mice compared to controls (**Figure 1H**). This variant morphology of *RPR2S* tumors was associated with markedly lower levels of CGRP, a well-described NE marker (**Figure 1I**). To determine if the reduction in CGRP resulted from loss of NE differentiation or if *RPR2S* tumors were initiated in a non-NE cell type, *RPR2* and *RPR2S* mice were infected with adenovirus Cre driven by the *Cgrp* promoter (Ad-CGRP-Cre) to induce tumorigenesis specifically in NE cells(19). After infection with Ad-CGRP-Cre, *RPR2S* tumors showed a significant reduction in levels of CGRP, confirming the loss of NE features is due to *Smarca4*-loss (**Supplementary Figure 1C**).

NE differentiation is tightly controlled by ASCL1, a basic helix-loop-helix transcription factor necessary for tumor development in *RPR2* mice(20). Because the SWI/SNF complex and ASCL1 functionally interact during neural development(21), we questioned whether SMARCA4 loss influences ASCL1 expression in tumors. ASCL1 transcript and protein levels were significantly decreased in *RPR2S* tumors compared to controls (**Figure 1J**). A causal relationship between the proteins in preSCs was shown as inhibition of SMARCA4 using FHD-286 significantly reduced ASCL1 expression in RT-qPCR and immunoblots (**Supplementary Figure 1D**). Taken together, these findings suggest that SMARCA4 loss results in inhibition of tumor development and loss of NE differentiation via ASCL1 loss, uncovering the role of the SMARCA4-ASCL1 axis in early-stage SCLC development.

A recent study suggests that the spontaneous loss of NE differentiation occurs in a SCLC GEMM in which tumor development is initiated by deletion of *Rb1* and *Trp53* and driven by a stable form of MYC (henceforth *RPM*)(22,23). We tested if *Smarca4* deletion differentially affects tumor development in *RPM* mice (henceforth RPMS) (**Figure 2A**). Ten weeks after Ad-CGRP-Cre infection, the tumor burden was significantly lower in the lungs of *RPMS* mice compared to *RPM* littermates (**Figure 2B**). Notably, the levels of pHH3 were not significantly different between the *RPM* and *RPMS* tumors, while the levels of cleaved CASP3 were significantly increased in *RPMS* tumors (**Figure 2C**). It was also noted that SMARCA4 expression was retained in some of the tumors in *RPMS* mice as shown in immunoblots (**Figure 2D**). Similar to the observations in *RPR2* mice, the *RPMS* tumors had a significant reduction in *Ascl1* transcripts measured by RT-qPCR (**Figure 2E**) that was associated with loss of CGRP (**Figure 2F**). These findings along with the observations in the *RPR2* model (**Figure 1I-J**) clearly indicate a crucial role of SMARCA4 in regulating ASCL1 and the tumor initiation capacity of SCLC. These results combined with the established necessity of ASCL1 for tumor formation in both the *RPR2* and *RPM* models(20,24) strongly suggest that the reduced tumor burden observed with loss of *Smarca4* is due, in part, to the disruption of the SMARCA4-ASCL1 axis. This relationship along with the retention of SMARCA4 in some *RPMS* tumors indicate its necessity for tumor initiation.

**Figure 2.**
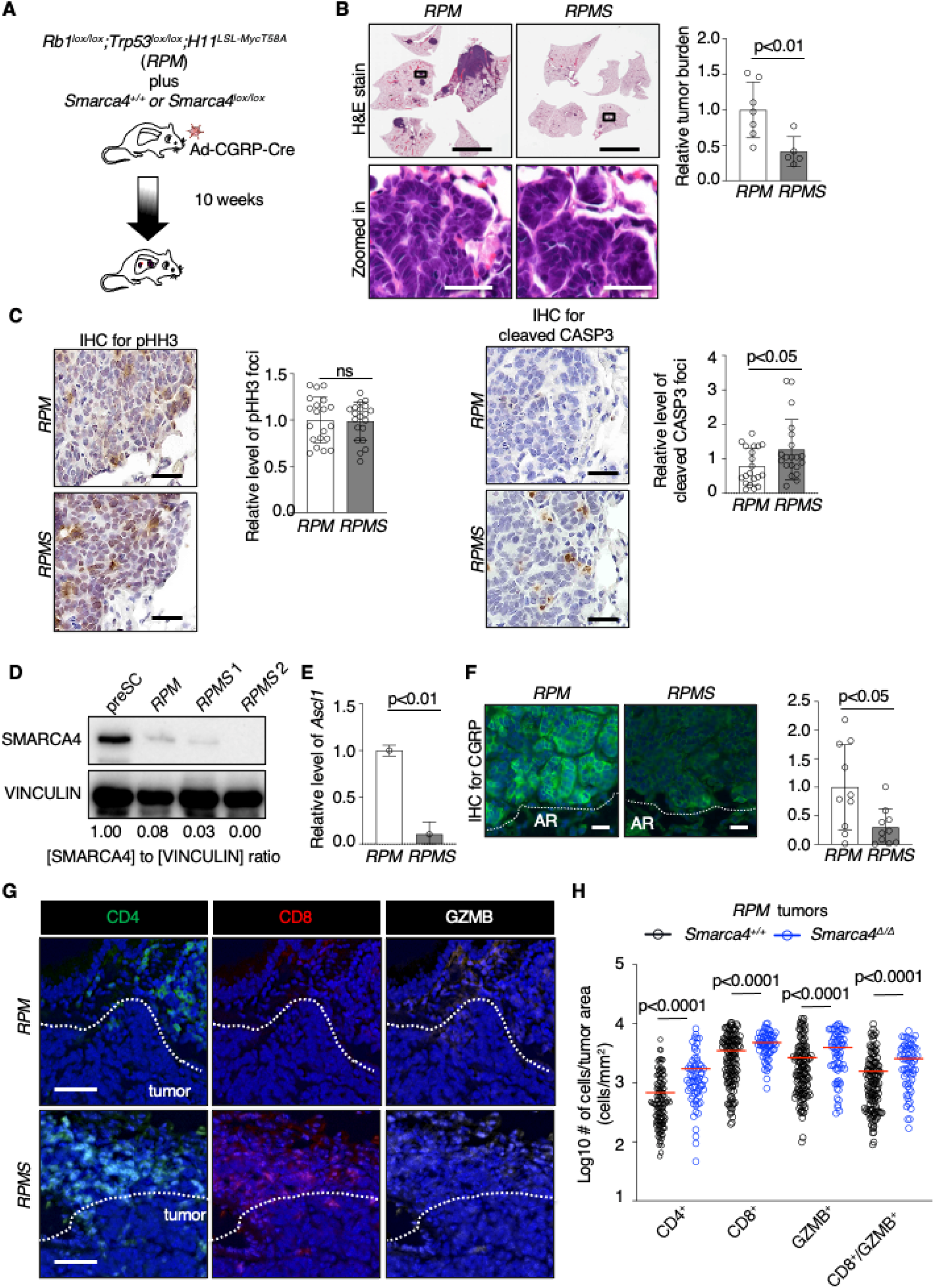
SMARCA4 is necessary for tumor development in *RPM* mice. **A**. Schematic of tumor induction in the *RPM* GEMM. **B**. Representative images of H&E-stained lungs of *RPM* (control) and *RPMS* (*Smarca4*-knockout) GEMMs, 10 weeks after Ad-CGRP-Cre infection (scale bars: 5mm) and zoomed-in images of specific tumor areas highlighted by the boxes in the images on top (scale bars: 50 µm). Right, quantification of tumor burden in the GEMMs normalized to the control *RPM* lungs (n=7 *RPM* and n=5 *RPMS*). **C**. Representative images of pHH3-and cleaved CASP3-stained *RPM* and *RPMS* lung sections (scale bars: 50 µm) and quantification of pHH3 foci and cleaved CASP3 foci within region of interest (ROI) in the lungs of *RPR2* and *RPR2S* (n=20 ROIs). **D**. Immunoblot for SMARCA4 expression in *RPMS* cells compared to preSC and *RPM* cells. Ratio of SMARCA4 levels relative to VINCULIN levels, measured by densitometry, shown on the bottom. **E**. RT-qPCR data showing *Ascl1* transcript level, normalized to levels of *Gapdh*, in *RPM* and *RPMS* cells (D) (n=3 per group). **F**. Representative images of CGRP-stained *RPM* and *RPMS* slides (scale bars: 50 µm) and quantification of CGRP levels (n=10 ROIs per group). AR: alveolar region. **G**. Representative images of multiplex IHC in *RPM* and *RPMS* slides stained for CD4, CD8, GZMB. Scale bars, 50 µm. **H**. Quantification of immune cells (CD4+ T cells, CD8+ T cells, GZMB+ cells, GZMB/CD8 Dual-positive cells) per tumor area (cells/mm^2^) in the lungs of *RPM* and *RPMS* mice. Each dot represents a nodular tumor (n=3 mice per group).

The decreased tumor burden in *RPR2S* mice may be attributed to the decreased proliferation and increased apoptosis observed in the tumors (**Figure 1C**). However, the *RPMS* tumors only displayed increased levels of apoptosis (**Figure 2C**). This led us to explore whether cell-extrinsic factors contributed to the slower tumor development in *RPMS* mice. The tumor microenvironment is complex and made up of various factors that can influence tumor growth. This includes immune cells that can hinder tumor growth and induce apoptosis. We performed multiplex IHC using the Vectra platform to visualize immune cell infiltration within *RPM* and *RPMS* tumors. SCLC is largely considered to be “immune-excluded” with immune cells aggregating around the tumor edge and rare infiltration into the core of patient tumors. (25). We observed a similar localization pattern of immune cells at the periphery of the tumors. The *RPMS* tumors and adjacent periphery had a significant increase in the number of CD4+ and CD8+ T cells (**Figure 2G-H**). CD8+ T cells can act directly on tumor cells through the release of Granzyme B (GZMB), a serine protease that can trigger apoptosis in target cells(26). The *RPMS* tumors had a significant increase in the number of CD8/GZMB dual-positive T cells (**Figure 2G-H**). These findings indicate an increase in both the number and activity of immune cells within the *RPMS* tumors (**Figure 2G-H**). We propose that both cell-intrinsic factors, including the disruption of the SMARCA4-ASCL1 axis, combined with the cell-extrinsic increased immune presence immediately surrounding the *RPMS* tumors contribute to the increased levels of apoptosis and ultimately the reduced tumor burden observed upon loss of *Smarca4*.

The most intriguing observation in *Smarca4*-deficient *RPR2* and *RPM* tumors is that the tumors, albeit smaller, display high-grade histology and molecular features of late-stage tumors, including loss of NE differentiation. These apparently conflicting phenotypes may be attributed to the shortcomings of the GEMMs in which *Rb1, Trp53* and *Smarca4* are simultaneously deleted - an event unlikely to occur in patient tumors. Additionally, we observed a spontaneous loss of SMARCA4 expression in more malignant models of SCLC progression (**Figure 2D**) and in a study showing gradual loss of NE features, defined by applying an established NE signature(27), in RPM cells during a 21 day period in culture (**Supplementary Figure 2**)(23), suggesting that the loss of SMARCA4 may occur during the later stages of tumor development. Analyzing the transcriptome data of SCLC PDXs, we noted that low *SMARCA4* expression correlated with higher levels of *MYC* (28,29)(**Figure 3A**) which was interesting given the decreased tumor burden in *RPMS* mice (**Figure 2B**). These observations led us to develop models that better recapitulated the temporal order of alterations in patient tumors. To determine the functional consequences of SMARCA4 loss in established tumor cells, we transduced *RPM* cells, that had maintained expression of SMARCA4 and NE status, with lentivirus containing either *Smarca4* or non-targeting shRNA (**Figure 3B**) and validated knockdown using immunoblots and RT-qPCR (**Figure 3C, Supplementary Figure 3A**). Doxycycline-induced *Smarca4* knockdown significantly reduced levels of *Ascl1* transcript and ASCL1 protein as assessed by RT-qPCR and immunoblots (**Figure 3C, Supplementary Figure 3B**). *Smarca4* knockdown did not significantly affect the colony-forming capacity of the *RPM* cells in soft agar assays (**Figure 3D**). This lack of impact is notable, compared to the inhibitory effect of SMARCA4 loss on the proliferation of preSCs and *Smarca4*-deficient *RPR2* tumor cells (**Figure 1**), and yet it does not support the role of SMARCA4 as a tumor suppressor. We reasoned that if not affecting cell-intrinsic changes *in vitro, Smarca4* knockdown could influence the tumorigenic capacity of tumor cells *in vivo*. To test this idea, we implanted 5x10^5^ cells in the flanks of athymic nude mice and measured tumor growth. *Smarca4* knockdown tumors grew significantly faster than controls (**Figure 3E, Supplementary Figure 3C**). Intriguingly, whereas levels of pHH3 was not significantly different between the groups, levels of cleaved CASP3 were lower in *Smarca4* knockdown tumors (**Supplementary Figure 3D**). We postulated that the enhanced tumor growth *in vivo* is due to altered tumor-host interactions, especially innate immune cells, such as NK cells, that remain functionally intact in athymic nude mice. Flow cytometry for NK cell interacting ligands on tumor cells showed that polio virus receptor (PVR/CD155) levels on pre-injected *Smarca4* knockdown cells were significantly reduced compared to controls (**Figure 3F, Supplementary Figure 4A**) while levels of other NK cell interacting ligands (PVRL2, ULBP1, RAE1, and CD276) were generally low and not significantly different between the groups (**Figure 3F, Supplementary Figure 4B**). RT-qPCR and immunoblots showed that PVR expression was significantly downregulated in *Smarca4* knockdown cells compared to controls (**Supplementary Figure 4C**). Additionally, analysis of chromatin immunoprecipitation sequencing (ChIP-seq) of four SCLC PDXs from a recent study revealed that SMARCA4 occupies regions of the *PVR* loci (**Figure 3G**)(8). Since PVR on tumor cells interacts with DNAX accessory molecule-1 (DNAM-1) on NK cell surface and can activate NK cell mediated cytotoxicity(30), the decreased PVR display may render *Smarca4* knockdown cells a selective advantage to avoid NK cell mediated killing in subcutaneous allografts.

**Figure 3.**
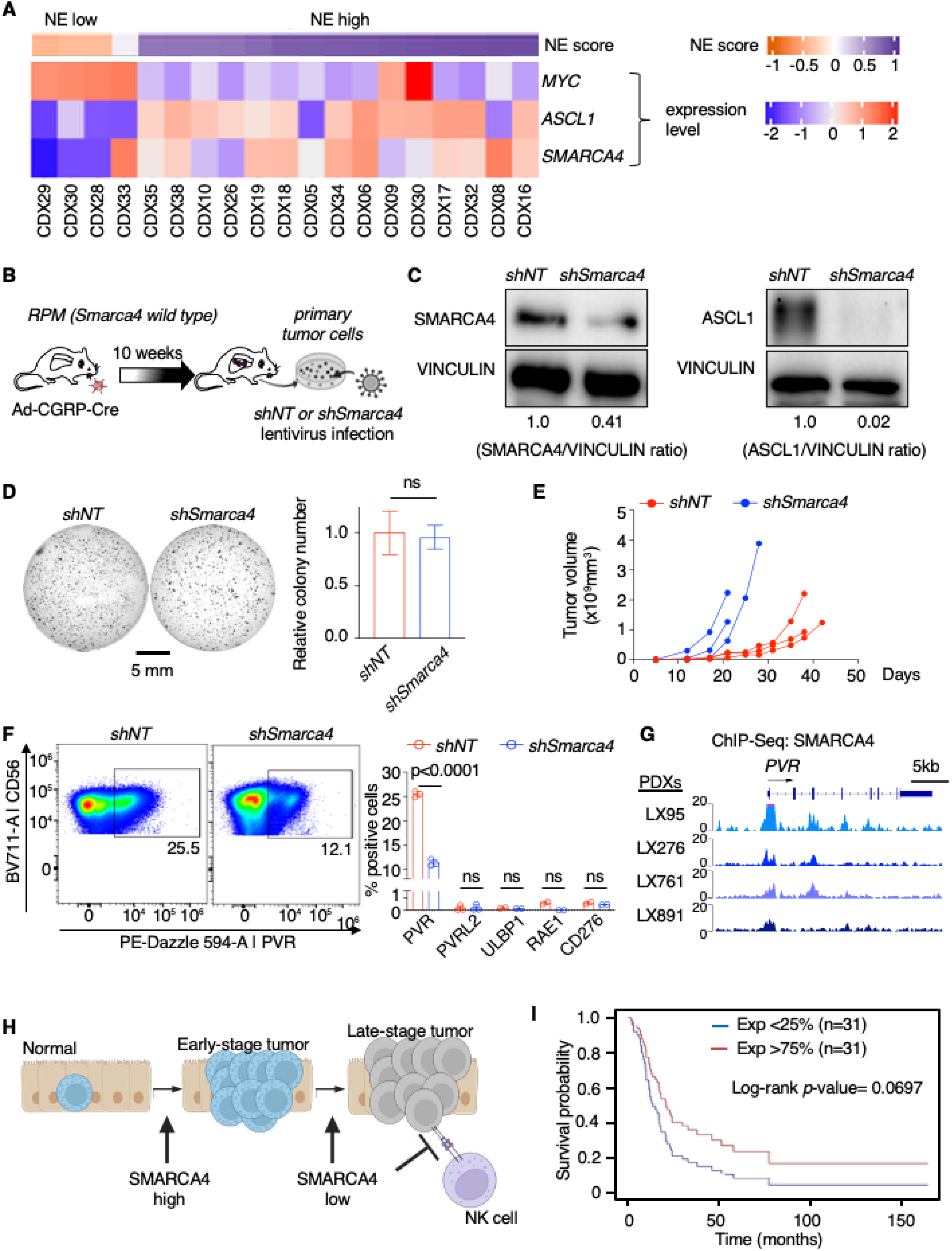
Acute SMARCA4 knockdown in tumor cells increased the tumorigenic potential. **A**. Heatmap showing expression of SMARCA4, ASCL1, MYC (rows) in 19 patient-derived xenografts (columns) arranged by NE score(27). Plot generated with the Gazdar Small Cell Lung Cancer Neuroendocrine Explorer(28). **B**. Schematic of targeting SMARCA4 in cells isolated from *RPM* tumors. **C**. Immunoblots for SMARCA4 and ASCL1 from non-targeted (*shNT)* and *shSmarca4* cells treated with 1 uM doxycycline for 48 hrs. Ratio of SMARCA4 levels to VINCULIN levels or ASCL1 levels to VINCULIN levels, measured by densitometry, shown on the bottom. **D**. Representative images of soft agar containing *shNT* and *shSmarca4 RPM* cells treated with 1uM doxycycline for the duration of the experiment and quantification of colony number. **E**. Measurements of tumor volume (mm^3^) of *shNT* and *shSmarca4* subcutaneous allografts in athymic nude mice (n=3 per group). **F**. Representative plots of flow cytometry quantification of PVR-positive cell population in *shNT* or *shSmarca4 RPM* cells treated with doxycycline for 48 hours. Right, flow cytometry quantification of positive PVR (n=3), PVRL2 (n=3), ULBP1 (n=2), RAE1 (n=2), and CD276 (n=2) cell populations in *shNT* and *shSmarca4 RPM* cells treated with doxycycline for 48 hours. **G**. ChIP-Seq for SMARCA4 at the PVR loci in 4 SCLC PDXs (LX95, LX276, LX761, LX891) adapted from (7). **H**. Model depicting the temporal-specific role of SMARCA4 in SCLC progression. **I**. Kaplan-Meier curve of SCLC patients from the highest and lowest quartile after stratification based on SMARCA4 expression (n=31 per group) (2,30). *P*-value by log-rank test is indicated.

Our findings provide novel insight into the temporal specific role of SMARCA4 in SCLC development. The SMARCA4 dependency in tumor development was discovered thanks to the approach of deleting *Smarca4* simultaneously with the tumor-initiating alterations in GEMMs. The expression pattern of SMARCA4 spontaneously decreasing from preneoplastic cells to tumor cells led to the idea that while required in preneoplastic lesions progressing towards tumor, it acts as a tumor suppressor in fully developed tumors (**Figure 3H**). The tumor suppressive effect of SMARCA4 is validated in part by our finding that acute *Smarca4* knockdown enhanced the tumorigenic capacity of tumor cells. To understand the impact of SMARCA4 levels in SCLC patients, patient tumors from two datasets(2,31) were stratified based on their SMARCA4 expression levels (either upper or lower quartile), revealed that low SMARCA4 expression correlated with worse overall survival (**Figure 3I**). This result is consistent with the tumor-promoting effect of low SMARCA4 levels observed in mouse models.

The concept of differential impacts of SMARCA4 with respect to the early versus late stage of SCLC is significant in that it addresses a fundamental question how the timing of tumor suppressor gene alterations shapes tumor evolution. However, the potential tumor suppressor role of SMARCA4 appears contradictory to its dependency in SCLC cells as suggested in recent studies where targeting SMARCA4 selectively inhibits the proliferation of POU2F3-positive SCLC cell lines representing a non-NE subtype of SCLC(7). The fact that the *Smarca4* knockdown *RPM* cells used in our study do not model the POU2F3-positive SCLC cells may explain the difference in SMARCA4 dependency. Along this line, another study showed that dual inhibition of SMARCA4/2 inhibited proliferation in patient, NE+ SCLC cell lines that are seemingly stuck in their differentiation state and are less likely to undergo loss of NE differentiation status, unlike the *RPM* cells used to knockdown *Smarca4*. While these recent studies showed the inhibitory effect of pharmacological dual-inhibition of SMARCA4 and SMARCA2 on SCLC cells, the effect could indicate a synthetic lethality among the components of SWI/SNF complex. It is also worth noting that *Smarca4*-knockdown enhanced the tumorigenic capacity of cells via a mechanism independent of cell proliferation. Our findings suggest a novel mechanism involving altered interactions between tumor and NK cells. Specifically, the down-regulation of PVR on *Smarca4*-knockdown cells reduces its binding to DNAM-1 to activate NK cell mediated cytotoxicity(30). This relationship is notable and requires further studies for functional validation. This SMARCA4-PVR axis may provide novel insight into the emerging role of NK cells in SCLC growth and progression(32,33) and suggest a need to evade NK cell surveillance in late-stage SCLC. In conclusion, the differential impact of losing SMARCA4 at the various stages of SCLC development revealed novel insights into the molecular patterns that drive the transformation of these tumors and reveal the varying vulnerabilities that may arise. SMARCA4 may prove to be an attractive target for SCLC prevention.

## Methods

This section is included in Supplementary Information.

