## Supplementary Information for "SMARCA4 is essential for early-stage tumor development but its loss promotes late-stage cancer progression in small-cell lung cancer"

**Materials and Methods**

*Mouse strains, Adenovirus Cre infection, and allografts*

The mouse strains employed *Rb1^lox/lox^; Trp53^lox/lox^; Rbl2^lox/lox^* (*RPR2*), *Rb1^lox/lox^;Trp53^lox/lox^* (*RP*), and *Rb1^lox/lox^;Trp53^lox/lox^; H11^lox-stop-lox-MYCT58A^* (*RPM*) and *Smarca4*^lox/lox^ were kindly provided by Drs. Anton Berns, Tyler Jacks, Julien Sage, Robert Wechsler-Reya, Trudy Oliver and Pierre Chambon and have been previously described(16,18,22,34). 6- to 8-week-old *RP* mice were intranasally administered with the adenovirus-CMV-Cre. For the RPR2 and RPM mice, adenovirus-Cre driven by either the *CMV* or *CGRP* promoter was intratracheally instilled in 10-week-old mice, including both male and female mice, for tumor induction. Ad-Cre viruses were purchased from the University of Iowa Gene Transfer Vector Core. Mice were aged 23 weeks for the *RPR2* model and 10 weeks for the *RPM* model. All mice were genotyped before and after experiments by polymerase chain reaction (PCR) of tail DNA which was purified using lysis buffer containing proteinase K (Thermo Fisher Scientific, BP100-100). Mice were maintained according to practices prescribed by the National Institutes of Health and all animal protocols were approved by the Institutional Animal Care and Use Committee (IACUC) at the University of Virginia. For the RP mice, all animal experiments were conducted in accordance with the regulatory standards approved by the Animal Ethics Committee of the Walter and Eliza Hall Institute of Medical Research (WEHI). For allograft experiments, 5.0 × 10^5^ murine cells were injected in the flanks of B6.129S F1 mice (the Jackson Laboratory). Following implantation, mice were fed a doxycycline diet (625 mg/kg, Envigo). Perpendicular tumor diameters were measured using calipers. Volume was calculated using the formula L × W2 × 0.52, where L is the longest dimension and W is the perpendicular dimension. The injected mice were maintained and observed for palpable tumors according to the procedures approved by the IACUC and euthanized when tumor size reached 1.5 cm in diameter, the endpoint of allograft study under the guideline of the institutional animal policy.

*Histology, Immunostaining, and Immunoblots*

At endpoint, the lungs were isolated and perfused with and incubated in 4% paraformaldehyde (PFA)/phosphate-buffered saline (PBS) for several days before paraffin embedding. Five micrometer-thick sections were stained with hematoxylin and eosin (H&E) staining. Additional slides were used for immunostaining. To dewax and hydrate the slides, they underwent sequential incubation in xylene, 100% ethanol, 95% ethanol, 80% ethanol, 70% ethanol, and PBS followed by antigen retrieval with EDTA. Antibodies used and their respective concentrations are listed in **Supplementary Table S1**. Macroscopic images of the lung sections were taken with the Olympus MVX10. Microscopic images of slides were acquired by the Nikon Eclipse Ni-U microscope. Image analysis and automated quantification of all H&E and immunostained slides completed using NIS-Elements Basic Research (Nikon). Quantification of immunostaining was performed on ≥5 randomly selected ROIs per tumor section. Thresholds were applied uniformly across samples. Scoring was conducted in a blinded fashion.

Immunoblots were performed with protein extracted from mouse tissue acquired from both tumor tissue and culture cells using RIPA buffer (50mmol/L Tris-HCl pH7.4, 150mmol/L NaCl, 2mmol/L EDTA, 1% NP-40, 0.1% SDS). Generally, 10 µg protein lysate was loaded into various percentages of SDS-polyacrylamide gels for electrophoresis in PAGE running buffer (25 mM Tris, 192 mM glycine, 0.1% SDS, pH8.3). Proteins were then transferred to 0.2 or 0.45 μm PVDF membrane in transfer buffer (25 mM Tris, 192 mM glycine, 20% methanol). Membranes were blocked for 1 hour at room temperature using TBST buffer (150 mM NaCl, 10 mM Tris pH8.0, 0.1% Tween20) supplemented with 5% non-fat milk, then blotted with primary antibodies overnight at 4°C on an orbital shaker. After washing 5 times with TBST, membranes were incubated with secondary antibodies for 2 hours at room temperature and washed another 5 times with TBST. Membranes were then incubated with ECL Western Blotting Detection Reagent (Thermo Fisher Scientific, 32106) for 5 minutes, and signals were detected using a ChemiDoc machine (Bio Rad). A list of all primary and secondary antibodies along with the concentrations utilized is listed in **Supplementary Table 1**.

*Multispectral Imaging*

Methods for staining of paraffin-embedded lung sections for multispectral imaging has previously been described(35) and were implemented(36). Antigen retrieval was performed using AR9 buffer (PerkinElmer) according to the manufacturer instructions. OPAL Multiplex IHC Staining (PerkinElmer) was performed according to the manufacturer instructions. Antibodies used and their respective concentrations are listed in **Supplementary Table S1**. Slides were mounted using prolong diamond antifade (Life Technologies) and scanned at low magnification using the Vectra 3.0 system and software (PerkinElmer). Regions of interest were identified in Phenochart software, and high-powered magnification images were acquired with the Vectra 3.0 system (Akoya Biosciences). These images were spectrally unmixed using single stain positive controls and analyzed using the InForm software (PerkinElmer). Immune cells were quantified using HALO software (Indica Labs, Albuquerque, NM).

*Flow Cytometry*

Cells were stained with a fixable live/dead cell stain (BioLegend Cat# 423105) at 1:1000 dilution for 15 min at room temperature, followed by staining with fluorophore-conjugated antibodies, listed in **Supplementary Table S1**, for 20 min on ice and in the dark. Cells were washed twice with a FACS buffer (1% BSA, 1mM EDTA in PBS) after every staining step and analyzed using an Aurora Borealis 5 laser Spectral Flow Cytometer at the University of Virginia Flow Cytometry Core, RRID: SCR_017829. Compensation was completed if samples were stained with multiple antibodies. The data were analyzed using FlowJo software. Example gating strategy is provided in **Supplementary Figure 4**.

*Quantitative RT-PCR*

For quantitative RT-PCR (RT-qPCR), total RNA was isolated from cells or tumors using the RNeasy Mini Kit (Qiagen, 74106) according to the manufacturer's protocol. cDNA was generated by the ProtoScript II First-Strand cDNA Synthesis Kit (New England Biolabs, E6560). RT-qPCR was performed using SYBR Green (Thermo Fisher Scientific, 4367659) with Applied Biosystems 7900 or StepOnePlus following the manufacturer's protocol. All qPCR primer sequences were retrieved from the online Universal Probe Library at Roche and included in **Supplementary Table S1**.

*Chemicals, vectors, and virus production*

A dual SMARCA2/SMARCA4 inhibitor, FHD-286, was obtained from MedChemExpress. Tet-pLKO-puro plasmid was obtained from Addgene (a gift from Dmitri Wiederschain). The shRNA oligo nucleotide sequences are listed in **Supplementary Table S1**. Lentivirus particles were produced by co-transfecting the Tet-PLKO-puro plasmid with packaging plasmids psPAX2 and pMD2.G (gifts from David Rekosh and Lou Hammarskjold), into HEK 293T/17 cells using polyethylenimine (Sigma-Aldrich, 408727). Forty-eight hours post-transfection, viral supernatants were harvested, centrifuged to remove cell debris, and stored at −80°C. Viral titers were determined by quantitative PCR (qPCR). Cells were transduced with lentiviral supernatant in the presence of DEAE-Dextran to enhance transduction efficiency. Puromycin selection (Thermo Fisher Scientific, A1113803) was applied 48 hours after transduction to enrich for successfully transduced cells, based on puromycin resistance conferred by the Tet-pLKO-puro vector. For inducible knockdown, puromycin-selected cells were treated with 1 uM doxycycline for 48 hours to induce shRNA expression. Knockdown efficiency was confirmed by RT-qPCR.

*Cells and soft agar assays*

Murine SCLC cells were isolated from lung tumors developed in the *RPR2* and *RPM* GEMMs. Precancerous cells (preSC) were derived from early-stage neuroendocrine lesions in the *RP* GEMM(17). All SCLC cells were cultured in RPMI-1640 media supplemented with 10% BGS and 1% penicillin-streptomycin-glutamine. 293T cells were cultured in DMEM supplemented with 10% BGS and 1% penicillin-streptomycin. For soft agar assays, 1x10^4^ cells per well in 0.5mL of growth medium containing 0.35% low-melting point agarose (Invitrogen, 16520-100_ and seeded on top of a 0.5mL base layer of medium containing 0.5% agar. After 3 weeks, colonies were fixed with 10% MeOH and 10% acetic acid and then stained with 1% MeOH, 1% Formaldehyde, and 0.05% crystal violet. Images of the wells were acquired using a Chemidoc and counted using the NIS-Elements Basic Research (Nikon) imaging software. All of the cell culture experiments were performed in triplicates and repeated for a minimum of two biological replicates.

*Gazdar Small Cell Lung Cancer Neuroendocrine Explorer*

The heatmap of RNA-seq Expression (log2 PRKM + 1) of SMARCA4, ASCL1, and MYC from 19 SCLC PDX samples(27) generated using the Gazdar Small Cell Lung Cancer Neuroendocrine Explorer (<https://lccl.shinyapps.io/GSNE/>) (28).

*Publicly available datasets (RNAseq, ChIP‑seq)*

RNA levels of SMARCA4 and NE markers in RPM cells cultured at varying timepoints were assessed from Ireland et al. (GSE149180)(22). Expression levels were downloaded and represented as FPKM (fragments per kilobase per million). The NE score was calculated using Zhang et al. signature(26). SMARCA4 ChIP-seq dataset was obtained from Redin et al. (GSE256346)(7).

*Statistical analysis*

GraphPad Prism was utilized for all statistical analysis as the mean ± standard deviation and evaluated using an unpaired two-tailed Student’s t-test. p<0.05 was considered statistically significant. Kaplan-Meier curves were used for plotting patient survival from (2,30) and log-rank test was used to determine significance with p<0.05 considered to be significant.

Supplementary Table 1. List of plasmids, oligonucleotides, and antibodies used

*Plasmids*

| **Plasmid** | **Source** |
| --- | --- |
| Tet-pLKO-puro | Addgene, 21915 |
| VSV-G | A gift from David Rekosh and Lou Hammarskjold at the University of Virginia |
| psPAX2 | A gift from David Rekosh and Lou Hammarskjold at the University of Virginia |

*shRNAs*

| **Target gene** | **Sequence (5' to 3')** |
| --- | --- |
| Non Target | ATCTCGCTTGGGCGAGAGTAAG |
| SMARCA4 | CGCCCGACACATTATTGAGAA |

*Genotyping primers*

| **Target alleles** | **Sequence (5' to 3')** |
| --- | --- |
| *Tp53+/lox* | Forward, CACAAAAACAGGTTAAACCCAG |
|  | Reverse, AGCACATAGGAGGCAGAGAC |
| *Rb1+/lox* | Forward CTCTAGATCCTCTCATTCTTCCC |
|  | Reverse, CCTTGACCATAGCCCAGCAC |
| *Rbl2+/lox* | Forward, GTGTTGTAACATTCTCGTGGG |
|  | Reverse, GACTGCTGGTATTAGAACCC |
| *Lox-stop-lox MycT58A* | Forward, AACTTCCCGCCGCCGTTGTT |
|  | Reverse, CAACGGGCCACAACTCCTCA |
| *H11 locus* | Forward, TGGAGGAGGACAAACTGGTCAC |
|  | Reverse, TTCCCTTTCTGCTTCATCTTGC |
| *Smarca4+/lox* | Forward, CATCTCACTCTTGTCCAGTC |
|  | Reverse, TCAGGGCAGACACAAAGT |

*RT-qPCR primers*

| **Target gene** | **Sequence (5' to 3')** |
| --- | --- |
| *Gapdh* | Forward, AGGTCGGTGTGAACGGATTTG |
|  | Reverse, TGTAGACCATGTAGTTGAGGTCA |
| *Smarca4* | Forward, GAAAGTGGCTCTGAAGAGGAGG |
|  | Reverse, TCCACCTCAGAGACATCATCGC |
| *Ascl1* | Forward, GCTCTCCTGGGAATGGACT |
|  | Reverse, CGTTGGCGAGAAACACTAAAG |

*Antibodies for Immunoblot and Immunostaining*

| **Target proteins** | **Source, Product Number, Analysis, Antibody Titer** |
| --- | --- |
| SMARCA4 | Cell Signaling Technology, 3508, Immunoblot, 1:1000, Immunostaining, 1:100 |
| ASCL1 | BD Pharmingen, 556604, Immunoblot, 1:750 |
| PVR | Abcam, ab103630, Immunoblot, 1:750 |
| VINCULIN | Santa Cruz, sc-73614, Immunoblot, 1:5000 |
| ACTB | Santa Cruz, sc-47778, Immunoblot, 1:5000 |
| CGRP (also known as CALCA) | Sigma-Aldrich, C8198, Immunostaining, 1:100 |
| phosphorylated histone H3 (pHH3) | Sigma-Aldrich, 06-570, Immunostaining, 1:100 |
| Cleaved CASP3 | Cell Signaling Technology, 9664, Immunoblot, 1:2000, Immunostaining, 1:100 |
| Secondary Ab-488 (Rabbit) | Thermo Fisher Scientific, A11034, Immunostaining, 1:200 |
| Secondary Ab-HRP (Mouse) | Jackson ImmunoResearch, 115-035-003, Immunoblot, 1:5000 |
| Secondary Ab-HRP (Rabbit) | Jackson ImmunoResearch, 111-035-003, Immunoblot, 1:1000 |

*Antibodies for Flow Cytometry*

| **Antibody** | **Dilution** | **Cat#** | **Vendor** |
| --- | --- | --- | --- |
| BV711 CD56 | 1:100 | 748099 | BioLegend |
| PE/Dazzle594 CD155 | 1:100 | 131516 | BioLegend |
| BV786 CD112 | 1:100 | 748050 | BioLegend |
| PE ULBP1 | 1:100 | FAB2588P | R&D Systems |
| PE RAE1 | 1:100 | 133203 | BioLegend |

*Antibodies for Multispectral Imaging*

| **Antigen Retrieval** | **Antibody** | **Dilution** | **Opal** | **Cat#** | **Clone** | **Vendor** |
| --- | --- | --- | --- | --- | --- | --- |
| AR9 | CD4 | 1:100 | 520 | ab183685 | EPR19514 | Abcam |
| AR9 | CD8 | 1:200 | 540 | ab237723 | CAL38 | Abcam |
| AR9 | Granzyme-B | 1:1000 | 690 | ab4059 |  | Abcam |

**Supplementary Figures**

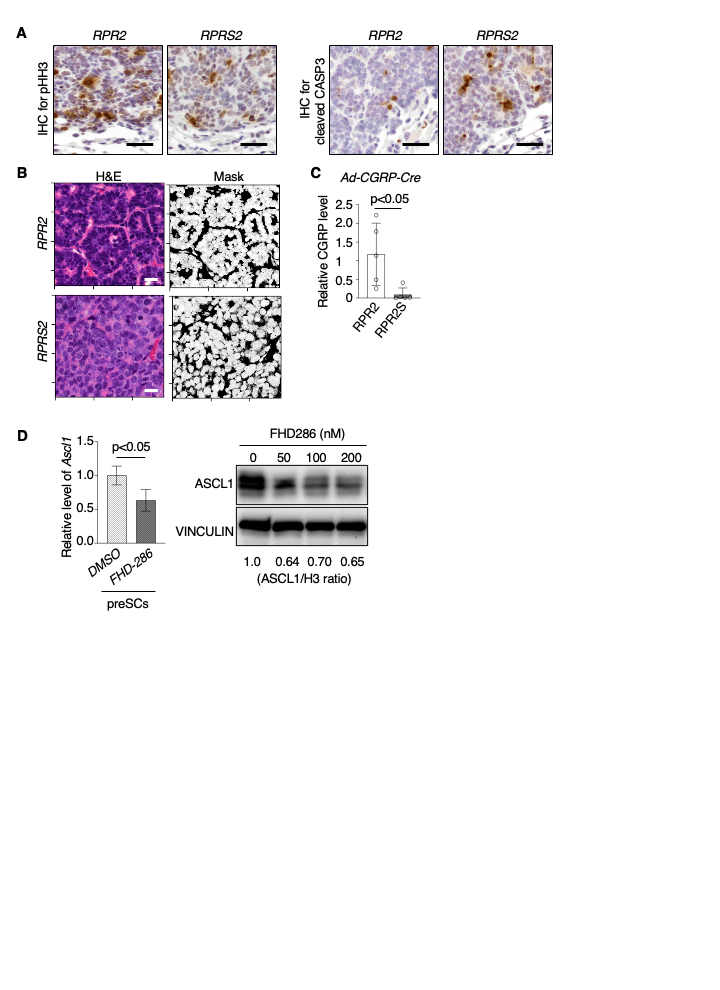

**Supplementary Figure 1. A.** Representative images of pHH3-stained or cleaved CASP3-stained *RPR2* and *RPR2S* slides. Scale bars: 50 µm. **B.** Left, representative H&E images used in CellProfiler classification. Scale bars: 50 µm. Right, CellProfiler generated masks of the H&E images. **C.** Quantification of CGRP expression in *RPR2* and *RPR2S* mice in which tumorigenesis was initiated using the Ad-CGRP-Cre virus, expressed relative to the *RPR2* controls, (n=5). **D.** Left, RT-qPCR data showing levels of *Ascl1* transcripts in preSCs treated with either DMSO or FHD-286 for 72 hours, (n=3). Data were normalized to levels of *Gapdh* transcripts and to the transcript levels in DMSO-treated preSCs. Right, immunoblots for ASCL1 and VINCULIN protein levels in DMSO (0 nM) and either 50, 100, or 200 nM FHD-286 treated preSCs for 72 hours. Quantification of protein levels shown at the bottom, expression normalized to levels of VINCULIN and to the protein levels in DMSO-treated cells.

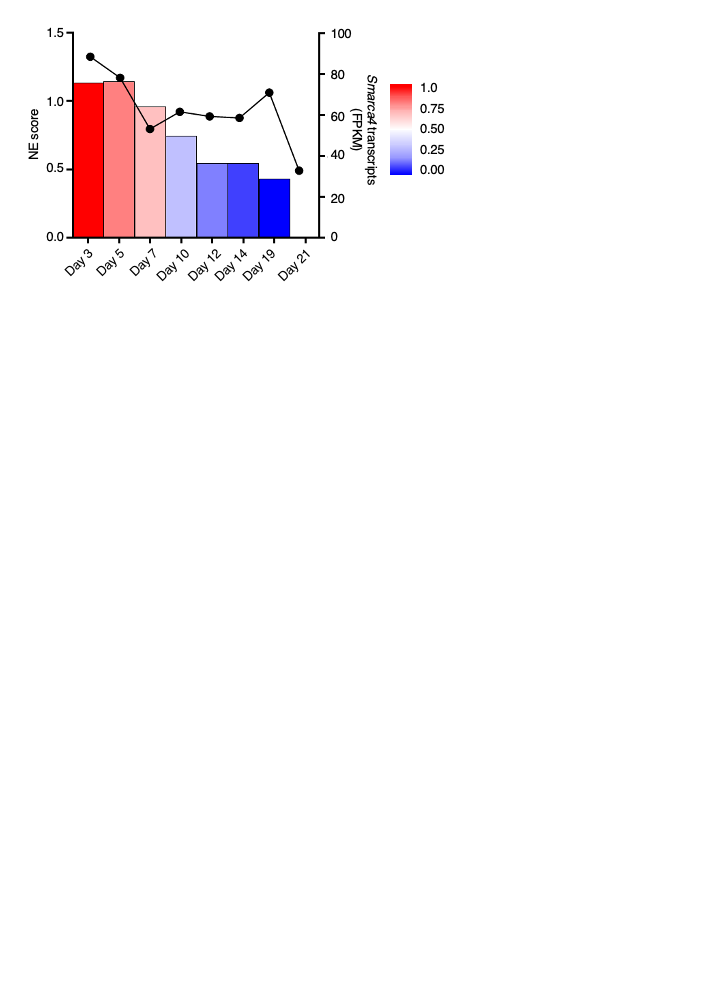

**Supplementary Figure 2.** Adapted from a study showing time course analysis of RPM cells undergoing loss of NE differentiation status(22). Expression of SMARCA4 shown as FPKM (fragments per kilobase per million) at the indicated timepoints relative to NE score(26).

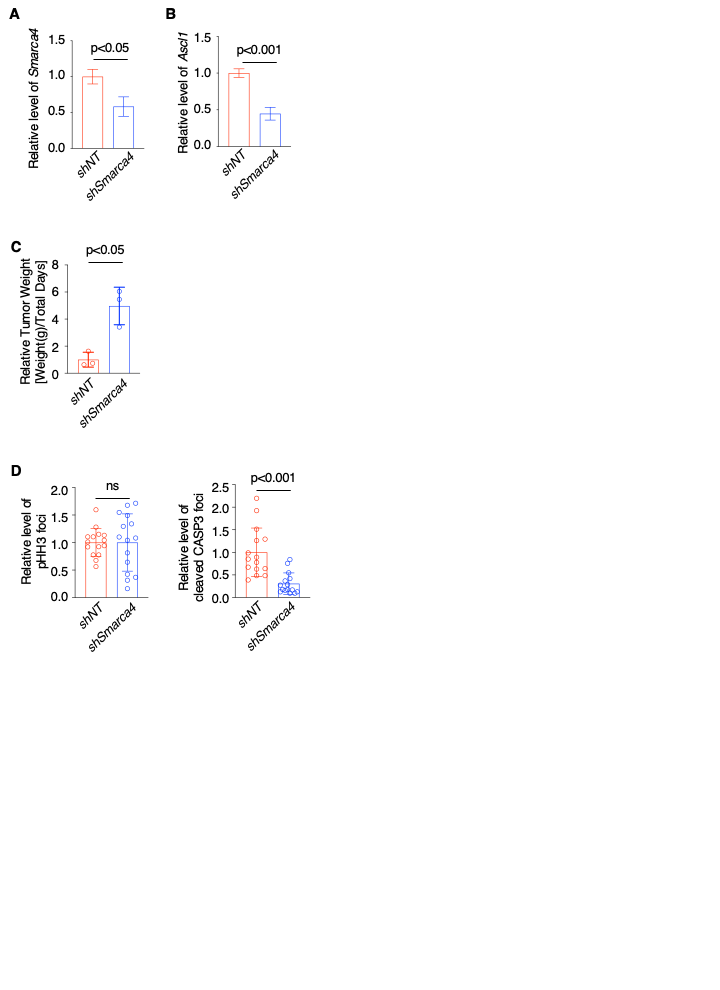

**Supplementary Figure 3. A and B.** RT-qPCR data showing levels of *Smarca4* and *Ascl1* transcripts in *shNT* or *shSMARCA4* *RPM* cells treated with 1 uM doxycycline for 48 hours (n=3). Data were normalized to levels of *Gapdh* and to the levels of transcripts in *shNT* *RPM* cells. **C.** Endpoint measurements of tumor weight expressed as the calculation of tumor weight (g) divided by the number of days post-injection and plotted relative to *shNT* controls. **D.** Quantification of pHH3 and cleaved CASP3 foci in *Smarca4* knockout tumors relative to the *Smarca4 wild-type* controls, n=15 ROI (5 ROIs per tumor).

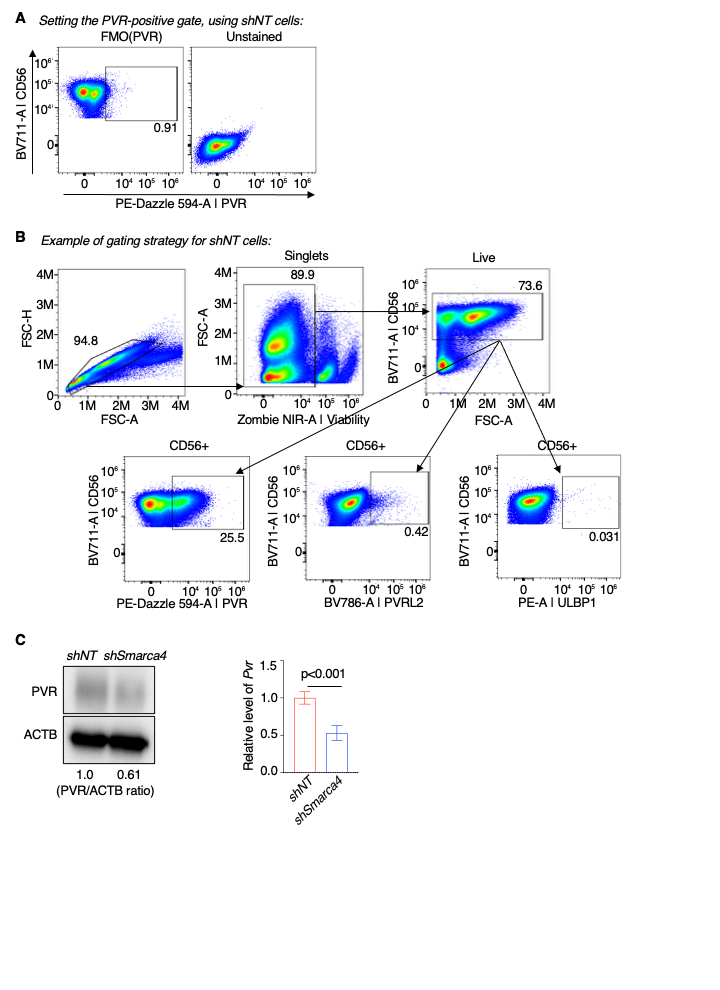

**Supplementary Figure 4. A.** Example of Fluorescence Minus One (FMO) gating strategy to define the PVR-positive population along with an unstained control. **B.** Example flow cytometry plots showing the progressive gating strategy, *shNT* sample displayed. **C.** Left, Immunoblots for PVR from *shNT* and *shSmarca4* cells treated with 1 uM doxycycline for 48 hrs. Ratio of PVR levels to ACTB levels, measured by densitometry, shown on the bottom. Right, RT-qPCR data showing levels of *Pvr* transcripts in *shNT* or *shSmarca4* *RPM* cells treated with doxycycline for 48 hours (n=3). Data were normalized to levels of *Gapdh* and to the levels of transcripts in *shNT* *RPM* cells.

**ACKNOWLEDGEMENTS**

This study was supported by NIH grants (U01CA224293 to K.S.P., R01CA278967 to J.I.P. and K.S.P; R01CA233661 to M.S.K., T32GM139787 to N.A.K), the UVA Parsons-Weber-Parsons Fellowship to N.A.K. A.T. was supported by the Center for Cancer Research, the Intramural Program of the NIH (ZIA BC 011793). This study was also supported by the NHMRC Project Grant (APP1159955) to K.D.S. This work was made possible through Victorian State Government Operational Support Program and the Australian Government NHMRC IRIISS. K.D.S is also generously supported by the Julie and Peter Alston Centenary Fellowship. The authors thank the Research Histology Core (RRID:SCR_025470), the Flow Cytometry Core (RRID: SCR_017829), and the Molecular Immunologic & Translational Sciences (MITS) Core at University of Virginia (NIH P30CA044579) and Leanne Scott and Ariena Kersbergen at the Walter and Eliza Hall Institute of Medical Research (WEHI) for their technical support and assistance of this work.

**CONFLICT OF INTEREST**

A.T. reports research funding from AstraZeneca, Tarveda, EMD Serono, and Prolynx. The rest of the authors do not have any conflict of interest related to this study.
